# Body Fluid Estimation during Standard Ultrafiltration in Chronic Kidney Disease

**DOI:** 10.1101/2025.04.15.648978

**Authors:** Rammah M Abohtyra, Omar A Beg

## Abstract

**Background:** Effective management of body fluid volumes and precise ultrafiltration (UF) prescription are critical challenges in treating Chronic Kidney Disease (CKD) patients undergoing hemodialysis (HD). Current fluid estimation techniques rely on fluid infusion or restricted UF protocols, which are difficult to implement consistently in daily clinical practice.

**Objective:** This work aims to evaluate whether current blood concentration measurement techniques can identify fluid and absolute blood volumes during regular HD treatments with standard ultrafiltration (UF) profiles (constant rates).

**Methods:** The proposed method is independent of any specific hematocrit sensor, UF rate, or volume infusion protocol. It utilizes modeling and prediction algorithms to quantify errors in fluid volume estimations.

**Results:** The method was tested on model-generated data from two patients under constant UF profiles. Extracellular (plasma and interstitial) fluid and absolute blood volumes were accurately estimated. In one case, specific blood volume dropped from 65 mL/kg to 61 mL/kg, while in the other, it remained above the critical threshold of 65 mL/kg.

**Conclusion:** This estimation algorithm can be easily integrated into existing HD machines, potentially improving treatment outcomes for CKD patients.

## I. Introduction

CHRONIC Kidney Disease (CKD) reduces the quality of life, raises mortality rates, and poses a significant economic burden [1]. Most advanced CKD patients rely on hemodialysis (HD) for treatment, typically receiving 3-4 sessions per week, each lasting 3-5 hours, during which excess fluid is removed through ultrafiltration (UF) [2]. However, both excessively high and low ultrafiltration rates (UFR) are linked to increased morbidity, with mortality rates reaching 15-20% in the first year of HD treatment [3]–[6].

Determining excess fluid volume by targeting a patient’s dry weight is essential for fluid management. Dry weight, the minimum weight at which a patient can leave HD treatment without symptoms like edema or intradialytic complications, is estimated only by clinicians [7]. Accurate estimation of dry weight reduces cardiovascular risks and mortality rates, while errors can lead to inappropriate therapy and severe outcomes [8], [9].

The standard UF protocol, typically involving a constant rate, is commonly used in daily practice but often employs high rates, resulting in an imbalance between fluid removal and interstitial-to-intravascular fluid refilling. [7], [10]. This results in a significant reduction in absolute blood volume, especially toward the end of HD therapy when fluid refilling slows. This imbalance can cause intradialytic complications such as hypotension, nausea, dizziness, and cramps, occurring in 25-60% of HD treatments [11].

Intradialytic hypotension (IDH), resulting from a significant reduction in absolute blood volume, is a common complication of HD and is linked to an increased risk of cardiovascular morbidity and mortality [12], [13]. Notably, an absolute blood volume threshold of 65 mL/kg has been associated with the onset of IDH events [14].

Direct measurement of absolute blood volume during HD is not feasible. Instead, relative blood volume (RBV) is estimated by monitoring changes in specific blood components such as hematocrit (H via Crit-Line monitor), hemoglobin (HGB), or total blood protein concentration (TBP). Several online measurement techniques are now available in clinical practice, but they are used only for estimating RBV changes during HD and UF [15], [16].

All current methods for measuring absolute blood volume rely on fluid infusion (indicator dilution) protocols, which require precise administration and measurement of the indicator, or they use restricted UF profiles. However, these techniques are not easily applicable for routine use [14], [17].

The aim of this study is to provide proof of concept for evaluating whether current techniques of measuring hematocrit, total protein, or hemoglobin concentration can also be used to identify fluid and absolute blood volumes during regular blood filtration treatment under constant UF profiles. This approach, proposed in this manuscript, does not rely on specific UF or infusion protocols. Instead, it fits a blood volume model to blood concentration data collected during a HD session in clinical practice. An optimization algorithm combined with a prediction algorithm, utilizing a modified two-compartment fluid volume model is used to achieve precise estimates of fluid and absolute blood volumes.

The paper is organized as follows: Section II outlines the estimation approach, Section III presents its application to blood filtration, Section IV presents the results, Section V is for discussion, and Section VI contains the conclusion. The Appendix is in Section VII.

## II. Materials and Methods

### A. Estimation Approach

The general form of our estimation approach utilizes a non-linear model of continuous-time differential equation described by a state transition (*f*) and observation function (*h*), given in this form

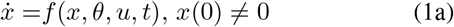

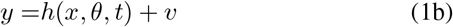

where *x ∈* ℝ^*q*^ is the unobserved state, *u* is the input, and *θ* ∈ ℝ^*p*^ contains model parameters, some of which are estimated from data; *y* is the output, *t*_0_ *≤ t ≤ t*_*f*_ is the time interval, and *v* represents measurement noise.

The estimation approach sequentially applies parameter and state estimation algorithms, as shown in Fig. 1. Estimated parameters are input to the unscented Kalman filter (UKF), which then estimates state variables using the nonlinear model (1) and accounts for errors from model uncertainty and measurement noise. The UKF is an extension of the Kalman filter that addresses the non-linearity in dynamic models more effectively [18].

**Figure 1.**
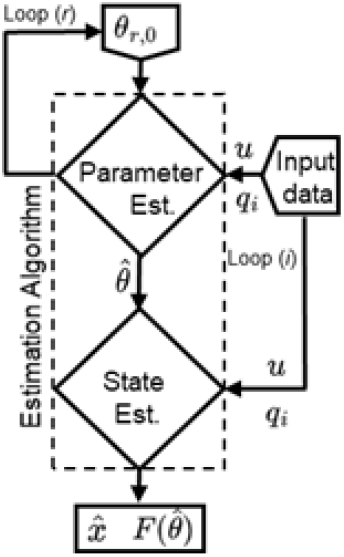
Parameter and state estimation approach (Algorithm 1). The parameter estimation and unscented Kalman filter (UKF) algorithms are separately performed. The estimated parameter 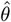 is an input to the UKF Algorithm.

#### 1) Parameter Estimation Algorithm

The algorithm optimizes the parameters *θ* by minimizing the sum of squared errors between the measured values (*z*_*j*_) and the model predictions (*y*(*θ, t*_*j*_)); *t*_*j*_ represents the time of the *z*_*j*_ measurements for *j* = 1, …, *N*. The objective function is given by

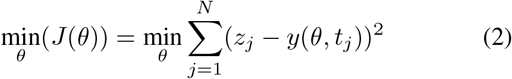

The optimal solution of (2) is denoted by 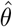. The algorithm has two nested loops (Fig. 1), the outer loop (*r*) iterates over random initial conditions *θ*_*r*,0_, while the inner loop (*i*) uses the Levenberg-Marquardt method to update *θ*_*r*_ with *θ*_*r,i*+1_ = *θ*_*r,i*_ *− ∇*_*r,i*_, where *∇*_*r,i*_ follows the steepest descent method [19].

Note that the estimated parameter 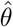 is an input to the UKF prediction algorithm.

#### 2) UKF Prediction Algorithm

To estimate parameter uncertainty and predict state trajectories using the UKF, we incorporate model uncertainty (*w*) into the dynamic equation:

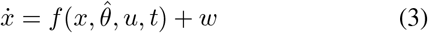

We assume that both model uncertainty (*w*) and measurement noise (*v*) in equation (1b) are piecewise constant over intervals of length *T*_*s*_ seconds (where *T*_*s*_ represents the sampling time). For each interval *k* (*k* is an index), these uncertainties are modeled as Gaussian random vectors with zero mean, described by

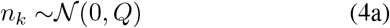

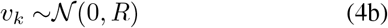

where 𝒩 denotes the Normal Gaussian distribution, and *Q* and *R* are the covariance matrices for model uncertainty and measurement noise, respectively. Thus, we assume that both the state *x* and the measurement *y*, which depends on *x*, follow Gaussian distributions. Consequently, the conditional state (*p*_*x*_) and output (*p*_*y*_) density functions are also Gaussian:

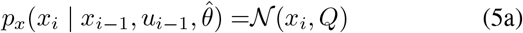

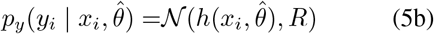

where *x*_*i*_ is the solution to (1) for *w* = 0, with *x*_*i−*1_ as the initial condition and *u*_*i−*1_ as the input.

As shown in Fig. 1, the algorithm first estimates 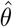 and then employs the UKF to compute the Fisher information matrix 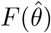, update the state, and predict the model output (1). The iterative steps for updating the UKF use observations *z*_0_, *z*_1_, …, *z*_*N*_ collected at sampling time *T*_*s*_, with the input *u* held constant within each interval (*i −* 1)*T*_*s*_ *≤ t ≤ iT*_*s*_:

- 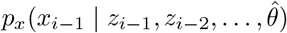 represents the probability of the previous estimated state.
- 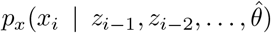 represents the prediction update step (time update), obtained by integrating model (1) between time *t* = (*i −* 1)*T*_*s*_ to *t* = *iT*_*s*_, with initial conditions 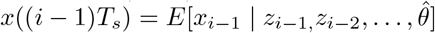.
- 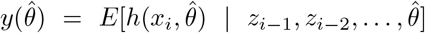 represents the prediction step of a measurement *z*_*i*_.
- 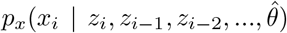 represents the measurement update step of the UKF.

Algorithm 1 integrates parameter estimation (A) with the UKF prediction algorithm (B).

##### Algorithm 1

Estimation algorithm

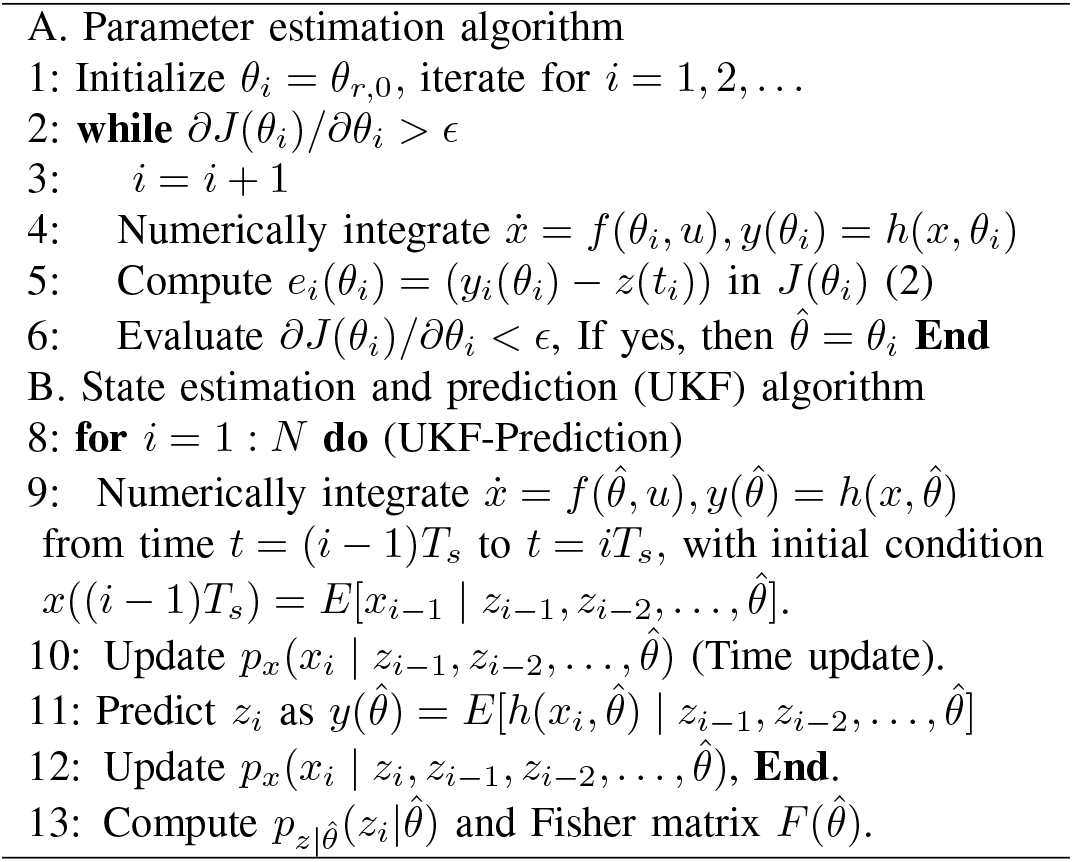

### B. Parameter Sensitivity Analysis

We employ the Fisher information matrix [20] to assess the quality of parameter estimation in the presence of model uncertainty and measurement noise. This analysis is based on the following assumptions.

- The distribution of *y* can we approximated by a Gaussian distribution.
- The experimental uncertainty is captured by model (1)
- The covariance of *y* is independent of 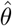.
- The UKF is approximately optimal, so that the prediction errors are uncorrelated.
- The parameter estimates are unbiased

#### Theorem 1.

*Under the assumptions listed above, the covariance of the parameter estimates (P*_*θ*_*) is bounded below by*

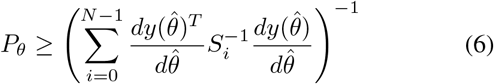

*where S*_*i*_ *is the covariance of the prediction error calculated using the UKF*.

**Proof:** For unbiased parameter estimates, the parameter estimate covariance is bounded below by the inverse of the Fisher information matrix (see e.g. [20]), which is calculated from the likelihood function. The likelihood function for the measurements *Z* where *Z* = [*z*_0_ *… z*_*N*_]^*T*^ is given by

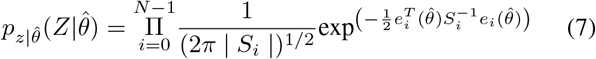

where 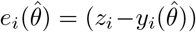, in which 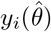 is the UKF one step ahead prediction; and *S*_*i*_ = *HP*_*i*_*H*^*T*^ + *R* is the covariance of the error 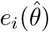, where *P*_*i*_ is the estimated state error covariance at time *i*, and *H* is the mapping from state to output (i.e., *H* = [1 0] for the model (8)). Then the log-likelihood function is

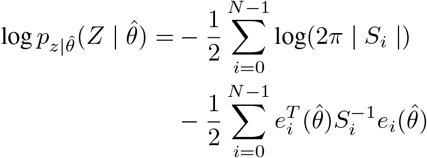

Then

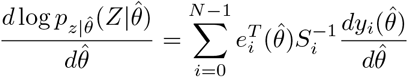

The Fisher information matrix is given by the covariance of this expression, so we find

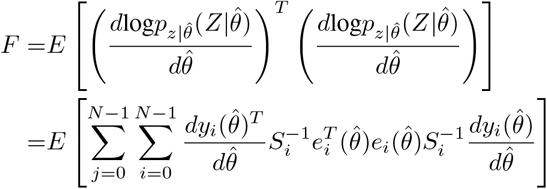

Under the assumption that the prediction error 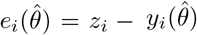 is uncorrelated,

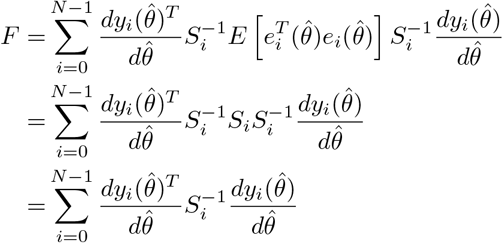

The result follows by inverting the Fisher information matrix. □

Throughout the paper, data are presented as mean *±* standard deviation (SD) where 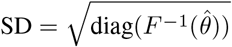.

## III. Application of Hemodialysis Treatment

In this application, as illustrated in Fig. 2, the proposed estimation approach leverages a model and blood concentration data from conventional HD sessions to estimate key parameters and body fluid volumes.

**Figure 2.**
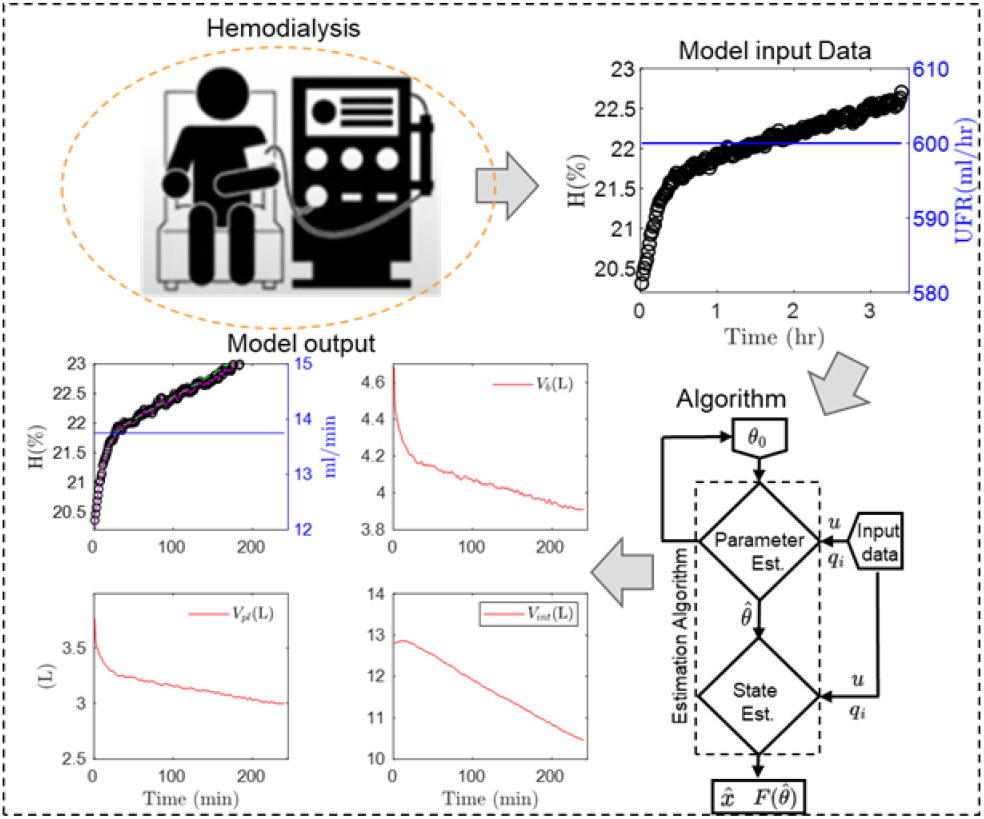
Body fluid estimation. Standard HD data (H and constant UFR) were used with the proposed algorithm to estimate plasma (*V*_*pl*_), interstitial (*V*_*int*_), and absolute blood (*V*_*b*_) volumes.

In our recent clinical study [21], the method was applied to dynamic hematocrit data from ten patient using two high-step UF profiles across 21 treatments and compared to an alternative method. However, this study demonstrates the feasibility of using the method in regular HD treatments with standard UF profiles, specifically constant UF profiles, and examines the estimated fluid and blood volumes under these profiles.

### A. Fluid Volume Model

The estimation algorithm employs a modified two-compartment extracellular fluid volume model to monitor volume kinetics during UF and estimate fluid volumes and parameters [22]–[25]. The model features two states: plasma volume (*V*_*pl*_ L) and interstitial fluid volume (*V*_*int*_ L). The ultrafiltration rate (*Q*_*u*_ L/min) serves as the model’s input, while the output is the whole-body hematocrit. Hematocrit (*H*) quantifies the volume fraction of red blood cells (*V*_*rbc*_) in relation to the total blood volume (*V*_*b*_ = *V*_*pl*_ + *V*_*rbc*_) and is expressed as *H* = *V*_*rbc*_*/V*_*b*_, typically represented as a percentage (%) in clinical settings.

The mathematical description of this model during UF is given by the nonlinear differential system:

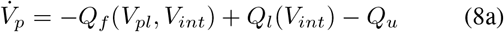

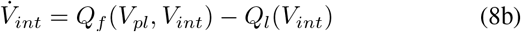

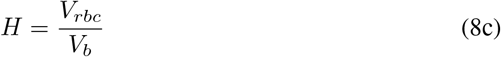

where *Q*_*f*_ (L/min) represents the microvascular filtration flow, encompassing both fluid shifts and refills. This flow is characterized by the transcapillary permeability coefficient, *K*_*f*_ (L/min.mmHg). Additionally, *Q*_*l*_ (L/min) indicates the lymphatic flow. Details of the model are explained in Appendix VII-A.

The model incorporates six patient-specific parameters, including initial plasma volume (*V*_*pl*_(0)) and interstitial fluid volume (*V*_*int*_(0)), represented as *θ* = [*V*_*pl*_(0), *V*_*int*_(0), *V*_*rbc*_(0), *K*_*f*_, *d*_1_, *d*_2_]^*T*^. Other parameters are assigned nominal values for all patients, as detailed in Appendix Table III.

## IV. Results

To evaluate the performance of this method under standard UF in a typical treatment setting, we generated model data using model (8), with parameters derived from the clinical study [21]. The model was driven by constant UF rates, utilizing parameters estimated from two patients undergoing variable UF, while integrating patient-specific details summarized in Table I, including, patient gender (G), pre-HD body mass (pre-M), post-HD body mass (post-M), UF volume (*V*_*u*_), and initial plasma portion concentration (*C*_*pl*_(0)), and more detail is found in [21].

**Table I.** PATIENT AND TREATMENT DESCRIPTION.

| Study | G | pre-M (kg) | post-M (kg) | $V_u$ (L) | $t_f$ (min) | $C_{pl}(0)$ (g/L) |
| --- | --- | --- | --- | --- | --- | --- |
| 1 | f | 53.5 | 50.5 | 3.3 | 240 | 20.3 |
| 2 | m | 62.2 | 58.5 | 5 | 300 | 21.6 |

In this illustrative example, the constant UF rates in the two studies were calculated by dividing the UF volume by the treatment duration (*V*_*u*_*/t*_*f*_), resulting in rates of 13.7 mL/min and 16.7 mL/min. Noisy hematocrit trajectories were generated by adding 10% random variation to each hematocrit value (Fig. 3 and 4).

**Figure 3.**
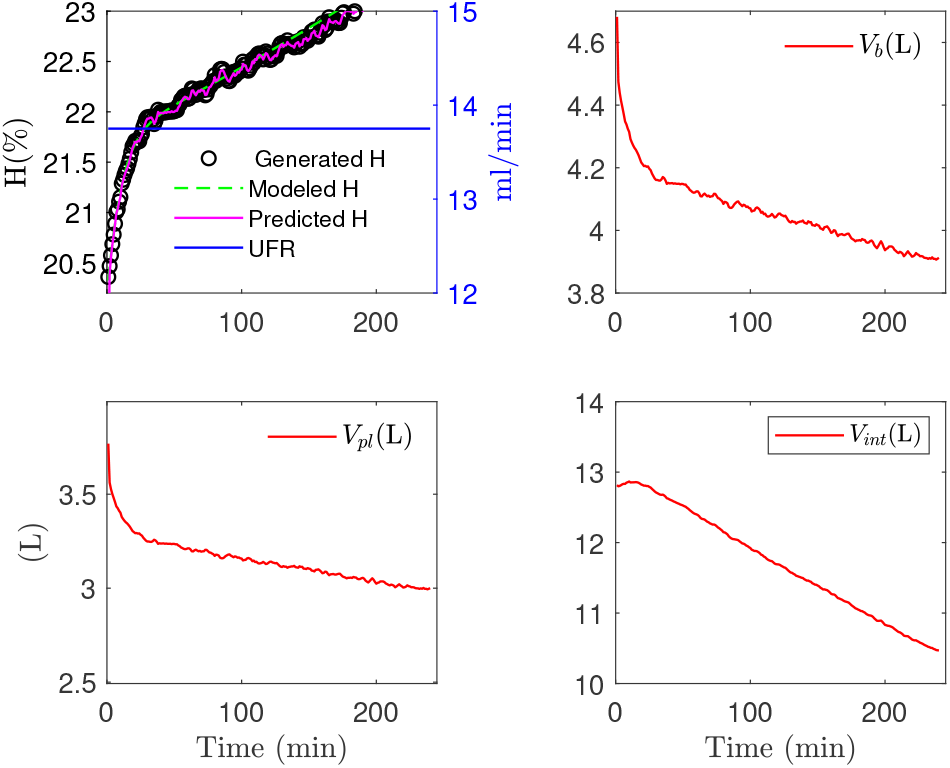
Modeling and estimation (Study 1). Top Left: Predicted hematocrit (H red) via UKF vs. modeled hematocrit (H green dashed) and generated data (H black) under constant UFR (blue, right). Top Right: Estimated absolute blood volume (red). Bottom Left: Estimated plasma volume (*V*_*pl*_). Bottom Right: Estimated interstitial volume (*V*_*int*_).

**Figure 4.**
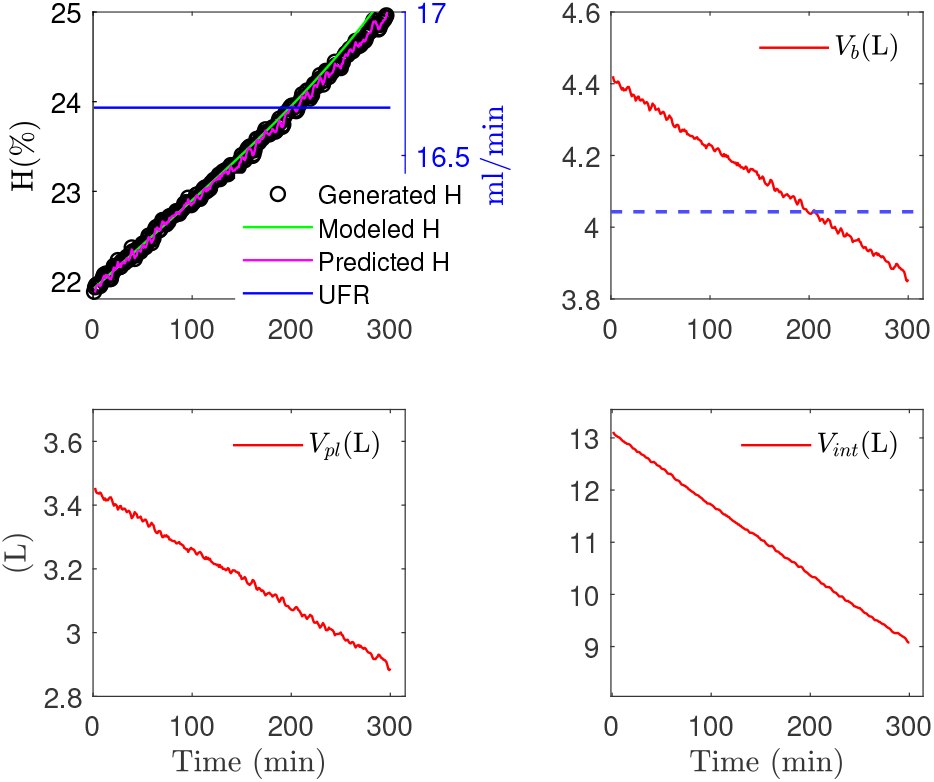
Modeling and estimation (Study 2). Top Left: Predicted hematocrit (H red) via UKF vs. modeled hematocrit (H green dashed) and generated data (H black) under constant UFR (blue). Top Right: Estimated absolute blood volume (red) and specific blood volume (65 mL/kg) occurs at 4.04 L (65 *×* 62.2/1000). Bottom Left: Estimated plasma volume. Bottom Right: Estimated interstitial volume.

A 30-minute segment of data, including hematocrit and UFR, was used to fit the model and estimate parameters, while the remaining data was reserved for validation.

The estimated parameters, presented in Table II, show an average error of just 5% compared to the generated data parameters, except for *d*_1_ and *d*_2_, which exhibit higher errors due to their insensitivity to hematocrit. The DS is calculated using the results from Theorem 1. The histograms (Fig. 6 and 5), generated from random initial conditions, show estimation distributions that are roughly normal and centered around the parameter maxima. The initial conditions were calculated by varying the patient-specific initial values by *±*50% [21].

**Table II.** MODEL GENERADED DATA PARAMETERS (*θ*) AND ESTIMATED PARAMETERS 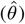.

| para. | study 1 |  | study 2 |  |
| --- | --- | --- | --- | --- |
| | model $\theta$ | estim. $\hat{\theta}$ | model $\theta$ | estim. $\hat{\theta}$ |
| $V_{pl}$ (L) | 3.5 | $3.35 \pm 1.6$ | 3.58 | $3.35 \pm 2.4$ |
| $V_{int}$ (L) | 13.21 | $13.42 \pm 5.7$ | 13.76 | $13.41 \pm 7.3$ |
| $V_{rbc}$ (L) | 0.98 | $0.93 \pm 0.13$ | 1.0 | $0.93 \pm 0.29$ |
| $V_b$ (L) | 4.5 | $4.7 \pm 1.7$ | 4.5 | $4.6 \pm 2.6$ |
| $K_f \times 10^{-3}$ | 6.4 | $4.7 \pm 4.4$ | 7.1 | $4.7 \pm 3.6$ |
| $d_1$ | 0.01 | $1.12 \pm 0.01$ | 0.13 | $1.12 \pm 0.07$ |
| $d_2$ | 0.07 | $0.02 \pm 0.07$ | 0.08 | $0.02 \pm 0.04$ |

**Figure 5.**
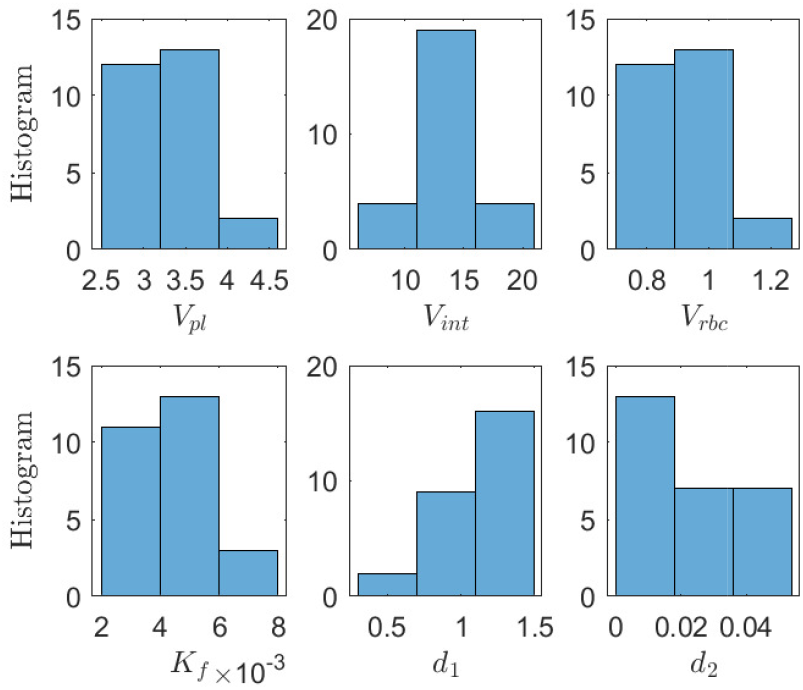
Histogram shows the sensitivity of the estimated parameters obtained using random initial values for Study 2.

**Figure 6.**
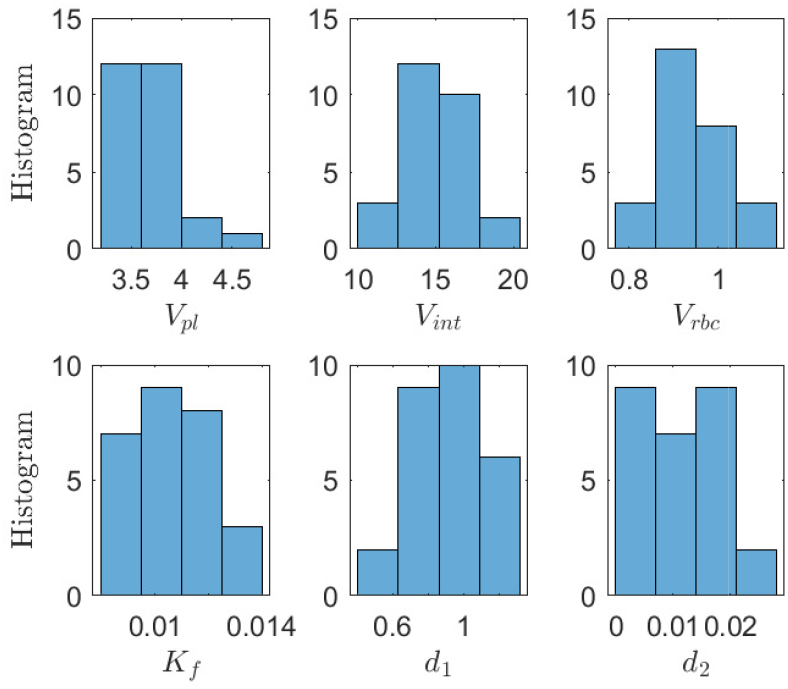
Histogram shows the sensitivity of the estimated parameters obtained using random initial values representing individual differences, for Study 1. The range (horizontal axis) reflects the accuracy of the estimates.

The predicted hematocrit trajectory (red line) using the UKF closely aligns with the generated hematocrit data (black circles). As shown in Fig. 3 and 4, the average error between the predicted and generated hematocrit is negligible, at 1.3 *×* 10^*−*4^ and 2.4 *×* 10^*−*4^, respectively. Similarly, the error between the modeled hematocrit (green dashed line) and the generated hematocrit remains small, at 1.3*×*10^*−*4^ and 1*×*10^*−*3^.

## V. Discussion

In this paper, we demonstrate the feasibility of our method by applying conventional constant UF profiles to model-generated data using a fluid volume model with parameters originally estimated from high-step UF profiles. Fluid and absolute blood volumes were predicted across two different studies. Our method enables the determination of specific blood volume thresholds (65 mL/kg) at which potential adverse events may occur.

As shown in Fig. 3 and 4, extensive constant UF leads to a significant decrease in both absolute blood volume and extracellular fluid volume by continuously removing plasma fluid. This reduction can potentially trigger IDH events. As shown in Fig. 4, at 203 min, the absolute blood volume drops to the specific absolute blood volume level of 65 mL/kg, requiring immediate interventions such as halting UF or administering volume substitution, both of which undermine effective volume management.

As shown in Fig. 3, during the first 30 minutes, the constant UF rate of 13.7 mL/min leads to a significant reduction in both fluid and absolute blood volumes. Afterward, an adequate refilling rate from interstitial space to intravascular space is observed, indicated by the rise in blood and fluid volume trajectories. During this treatment section, the absolute blood volume remains above the specific absolute value (65mL/kg). In this study, the specific absolute blood volume was 71.96 mL/kg at the end of the treatment section.

The high UF rate (16.7 mL/min) caused a continuous decline in both blood and fluid volumes, as indicated by a significant rise in the hematocrit trajectory (Fig. 4). In this study, the specific blood volume was at 4 L (65 *×* 62.2/1000), and by the end of the treatment, it had dropped to 62 mL/kg. We observed that when there is a good match between the modeled and generated hematocrit trajectories, the histogram of the estimated parameters (Fig. 6 and 5) from random initial values follows an approximately normal distribution, indicating high estimation quality. In contrast, poor modeling is reflected by a growing discrepancy between the modeled and generated hematocrit values, resulting in a wider, bimodal histogram.

The method can be fully automated without modifying the HD device hardware, but it is limited to responding to volume changes. Future work could focus on implementing automated feedback systems for blood volume-controlled UF to prevent drops below critical thresholds.

## VI. Conclusions

The proposed estimation approach offers a practical method for estimating body fluid volumes in regular blood filtration treatment, utilizing continuous hematocrit and ultrafiltration data without being limited by specific ultrafiltration rates or infusion protocols. Estimating absolute blood volume at the start and throughout each HD treatment session provides an opportunity to improve fluid management in CKD patients. This method can also identify critical blood volumes during individual HD treatments and detect low absolute blood volumes in advance. Its integration could support automated feedback systems to adjust and prevent drops below critical volume thresholds.

## VII. APPENDIX

### A. Appendix A:Fluid Volume Model

The mathematical description of *Q*_*f*_ and *Q*_*l*_ are:

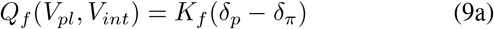

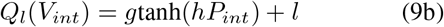

where *δ*_*p*_ and *δ*_*π*_ represent the hydrostatic and osmotic pressure gradients, respectively:

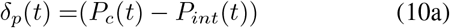

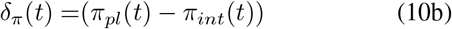

where *P*_*c*_, *P*_*int*_, *π*_*pl*_, *π*_*int*_ refer to the hydrostatic capillary pressure, interstitial pressure, plasma colloid osmotic pressure, and interstitial colloid osmotic pressure, respectively; and *g, h*, and *l* are constants (Table III). The equations for *P*_*c*_ (hydrostatic capillary pressure), *P*_*int*_ (interstitial pressure), *π*_*pl*_ (plasma colloid osmotic pressure), *π*_*int*_ (interstitial colloid osmotic pressure) in (10b) are given by

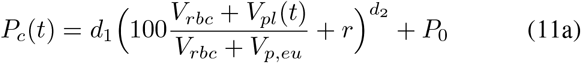

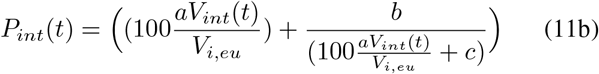

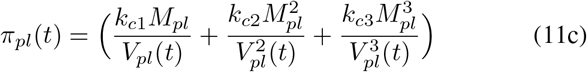

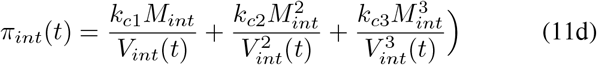

where *V*_*p,eu*_, *V*_*i,eu*_ refer to euhydration (normal) plasma and interstitial volume for 70 kg patient, respectively; *P*_0_ is an offset pressure; *M*_*pl*_ = *C*_*pl*_(0) × *V*_*pl*_(0) is plasma protein mass; *M*_*int*_ = *C*_*int*_(0) × *V*_*int*_(0) is interstitial protein mass; and *a, b, c, d*_1_, *d*_2_, *r, k*_*c*1_, *k*_*c*2_, *k*_*c*3_ are listed in Table III, as nominal parameters.

**Table III.** NOMINAL VALUES OF THE MODEL.

| parameters | value | units |
| --- | --- | --- |
| $a$ | 0.006 | unit-less |
| $b$ | -198 | unit-less |
| $c$ | -45 | unit-less |
| $r$ | -30 | unit-less |
| $g$ | 0.045 | unit-less |
| $h$ | 0.767 | unit-less |
| $g$ | 0.045 | unit-less |
| $l$ | 0.045 | unit-less |
| $P_o$ | 13.128 | (mmHg) |
| $V_{i,eu}$ | 11 | (L) |
| $V_{p,eu}$ | 3 | (L) |
| $k_{c1}$ | 0.21 | unit-less |
| $k_{c2}$ | 0.0016 | unit-less |
| $k_{c3}$ | $9 \times 10^{-6}$ | unit-less |

